## Supplementary Information for "Evaluating the predictive power of combined gene expression dynamics from single cells on antibiotic survival"

1. Supplementary Figures
2. Supplementary Tables
3. Supplementary Movie Captions

### Supplementary Figures

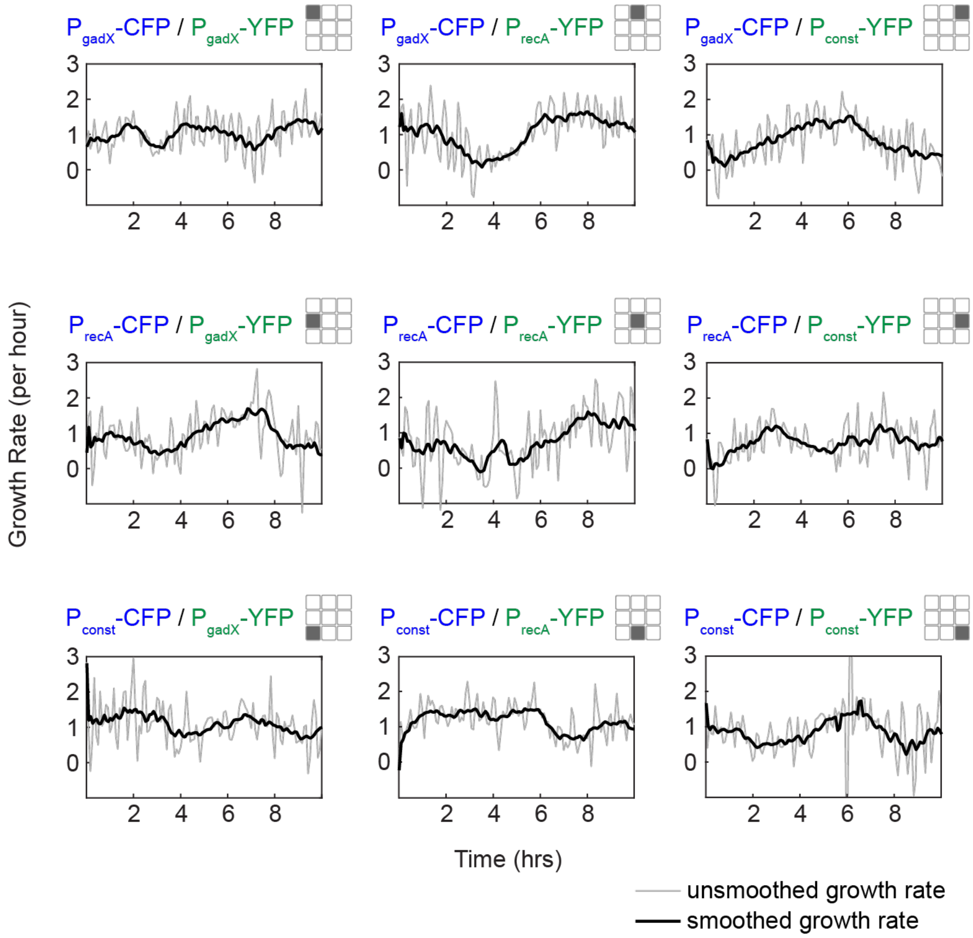


**Figure S1. Smoothing growth rate.** Growth rates of representative cells. Raw growth rate values extracted from the image analysis process are shown with a thin grey line, and smoothed growth rate with a 1 hour window are shown in a thick black line.

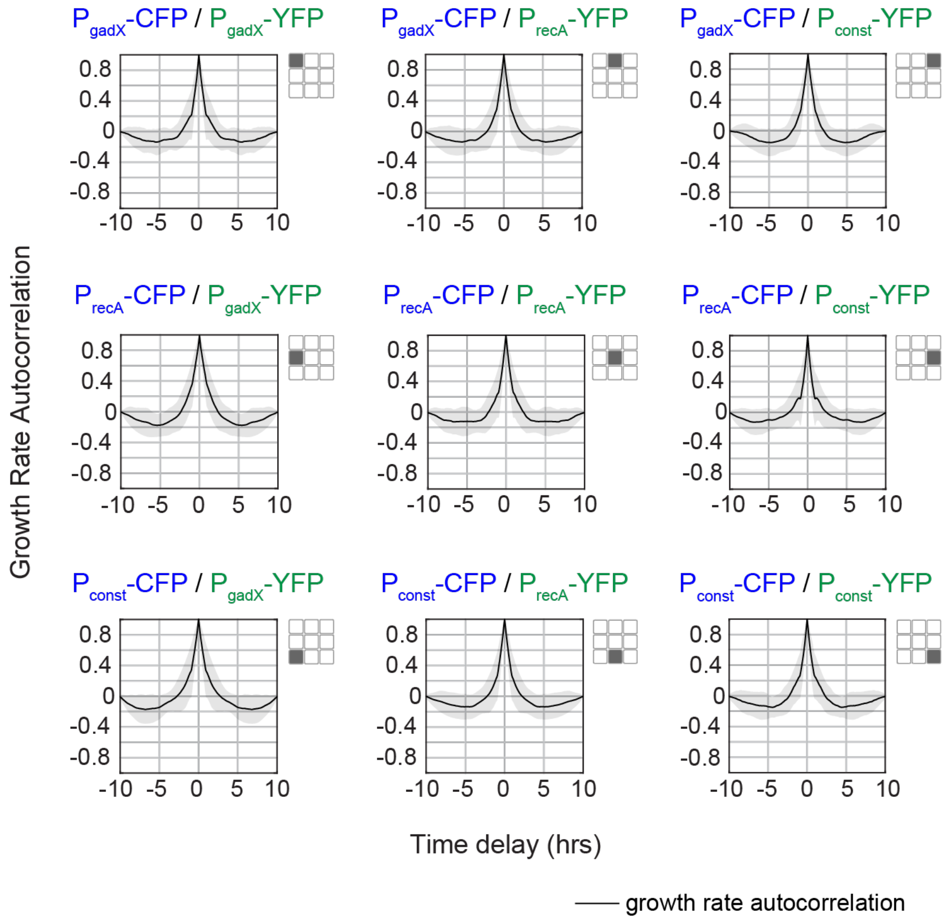


**Figure S2. Growth rate autocorrelations.** Autocorrelation of growth rates for each of the dual reporter strains.

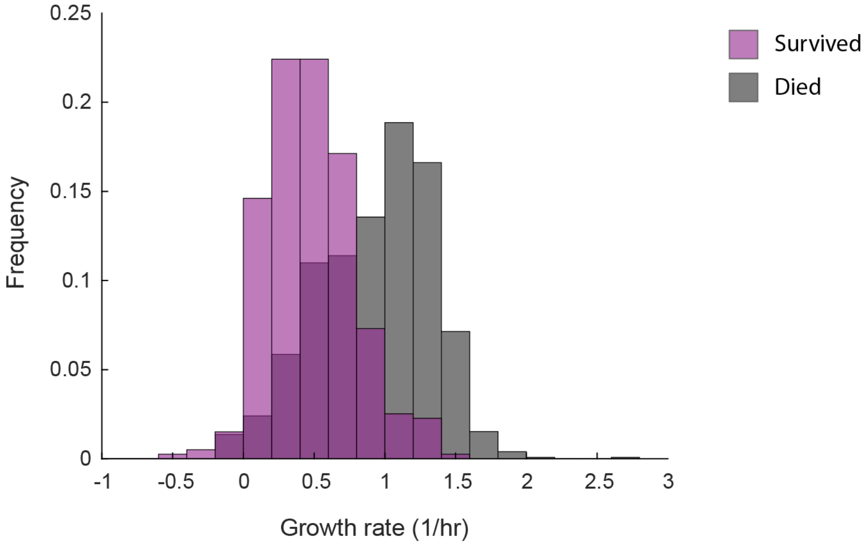


**Figure S3. Growth rate and ciprofloxacin survival.** Histogram showing the growth rate of cells that survived (purple) versus died (grey) after ciprofloxacin exposure.

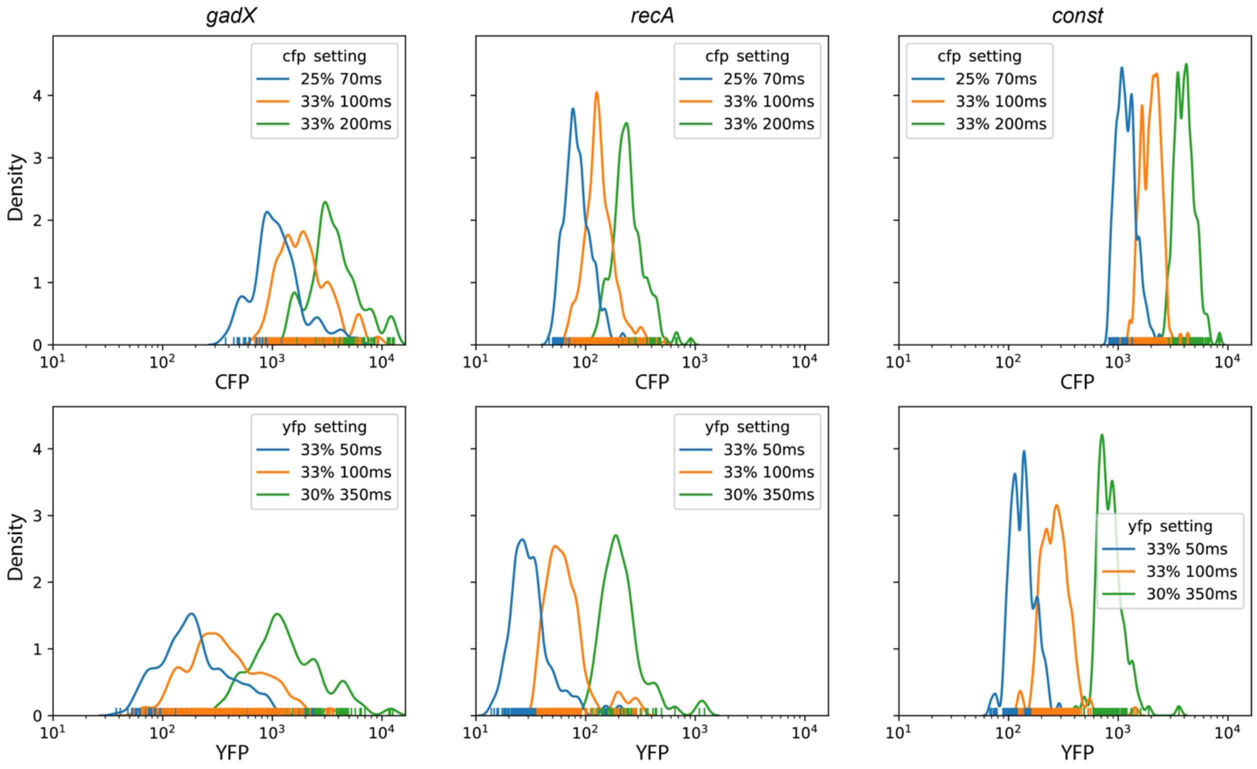


**Figure S4. Effect of the imaging setting on the fluorescence population distributions.** In calibration experiments, we imaged the same cell population with six different imaging settings, three for each color. We found that the fluorescence distributions translate without changing shape on a log scale, which indicates that the effect of the imaging setting is a multiplicative factor.

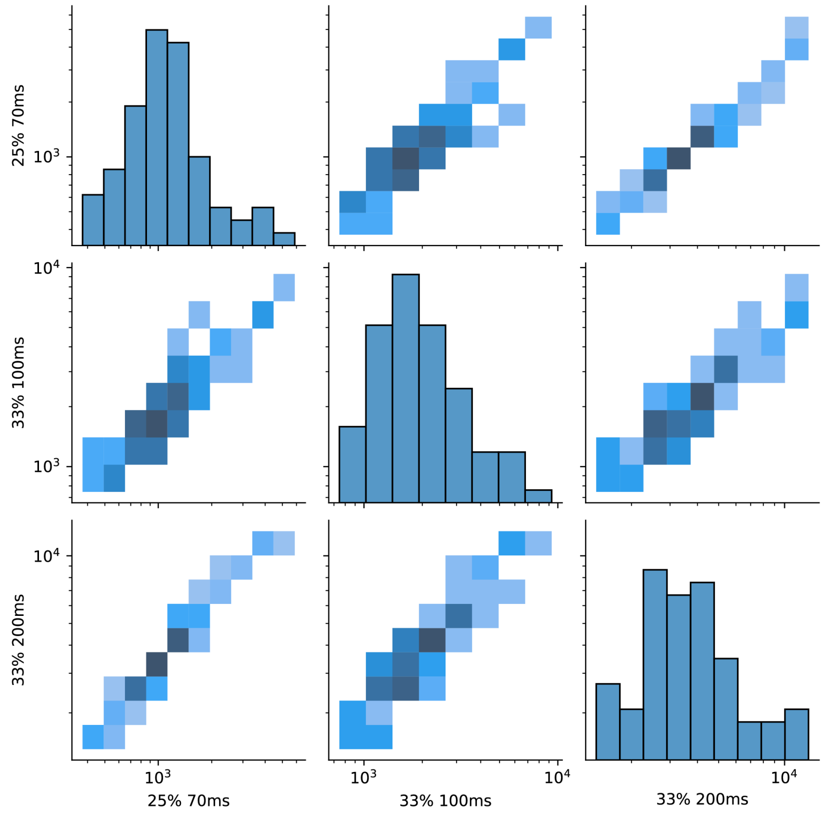


**Figure S5. Effect of the imaging setting on single cell fluorescence.** One-dimensional and two-dimensional histograms of the raw fluorescence values for $P_{gadX}$-CFP from calibration experiments. The presence of diagonal lines shows that the values across settings are largely proportional, which supports the multiplicative hypothesis even at a single-cell level. Other promoters ($P_{recA}$, $P_{const}$) and colors (YFP) are similar to this graph.

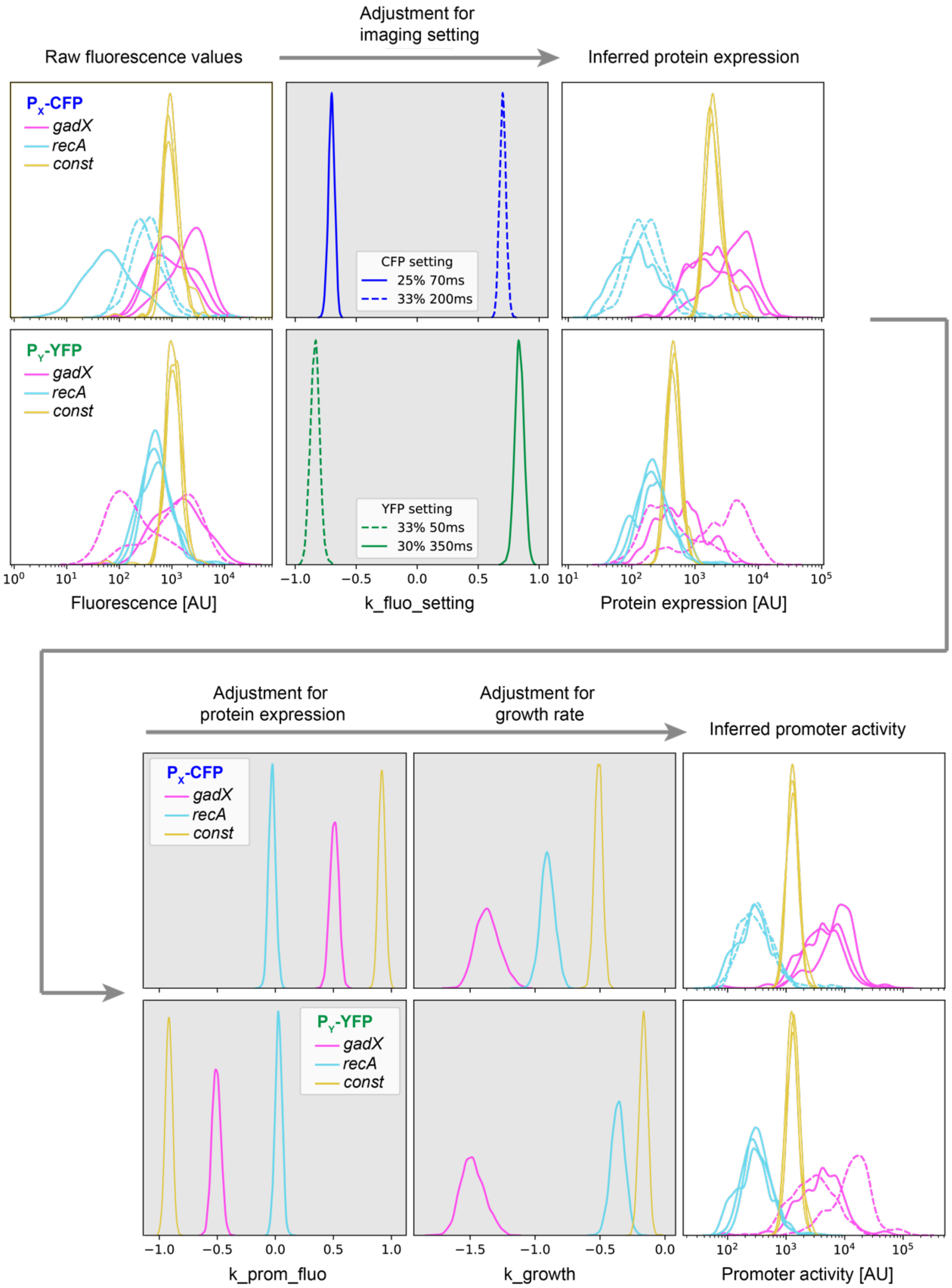


**Figure S6. Transformation of raw fluorescence values to inferred promoter activity.** This figure expands on Fig. 3C to show the posterior distributions of the parameters linking the raw fluorescence values, inferred protein expression, and inferred promoter activity. Refer to Methods for the details of the model.

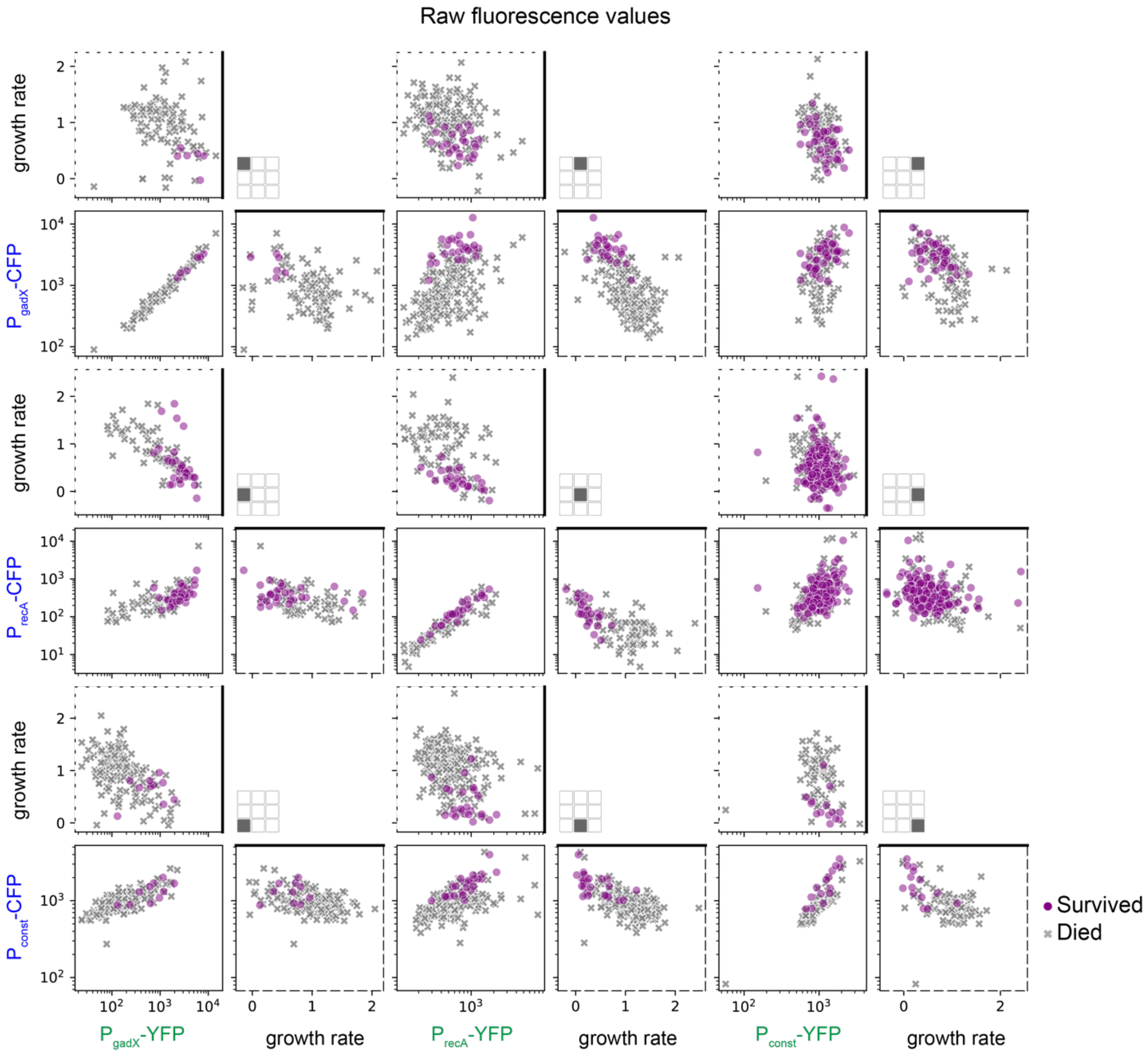


**Figure S7. Relation between growth rate, raw fluorescence values, and survival.** Raw data associated with Fig. 3E, before applying the corrections. Growth rate units 1/h; fluorescence units AU.

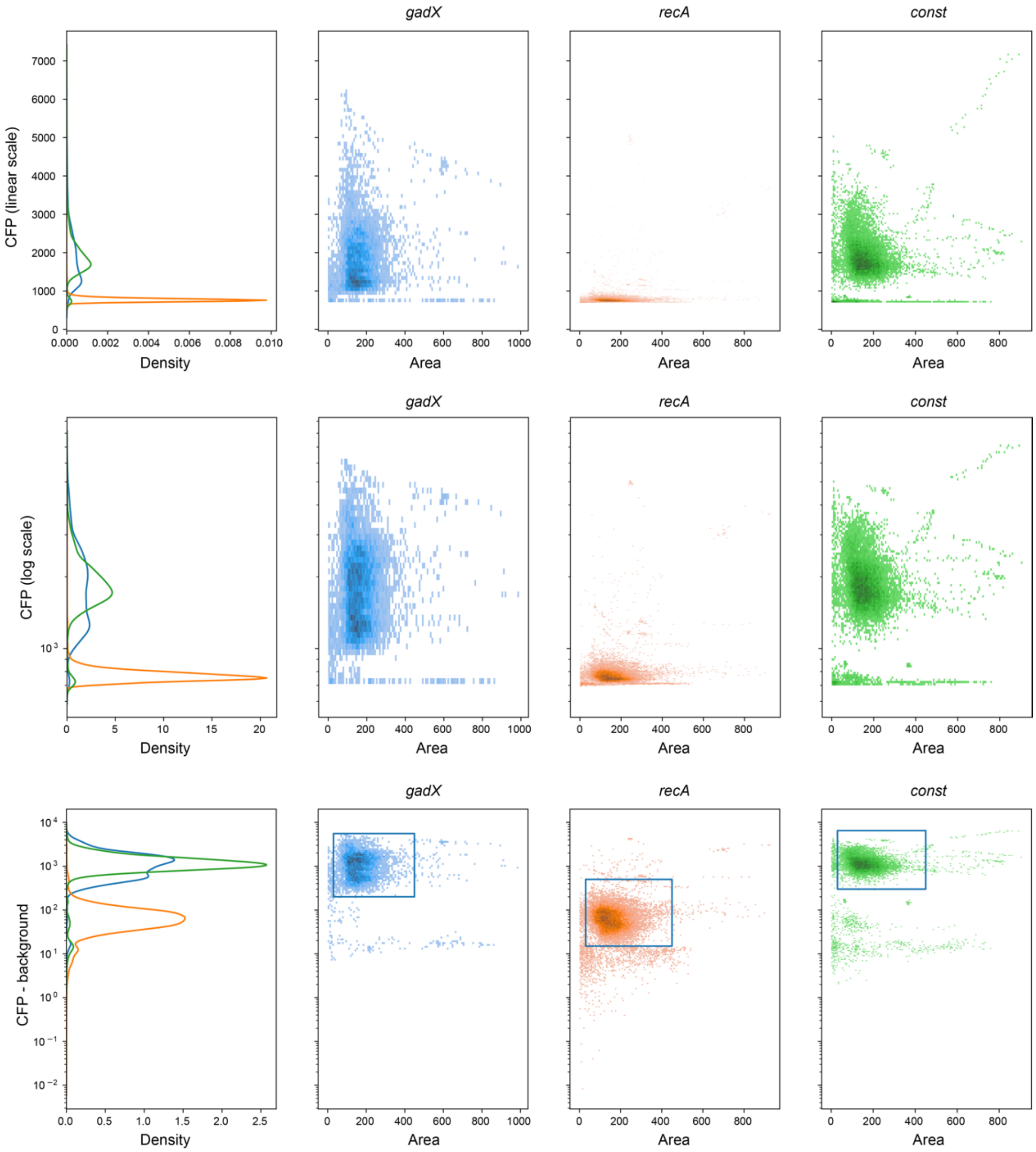


**Figure S8. After background subtraction, logarithmic scale is the natural scale for fluorescence distributions.** In the first row, raw fluorescence distributions as a function of cell area on a linear scale. Second row, the same data plotted with a logarithmic scale for fluorescence. Third row, the scale is logarithmic after we remove the background fluorescence. On every row, the first column is the marginal distribution of fluorescence. This figure shows the same data represented in three different ways, but illustrates that the third representation is more regular than the first two. The boxes indicate the gating values, which set the points that we retained and those that we did not. Data shown here are for $P_{gadX}$-CFP, $P_{recA}$-CFP, and $P_{const}$-CFP using the 25%-70ms setting, and are representative of all data.

### Supplementary Tables

**Table S1.** Plasmids used in this study.

| $P_x$-CFP | $P_y$-YFP | Origin | Resistance Marker | Addgene Plasmid ID |
|---|---|---|---|---|
| $P_{gadX}$-CFP | $P_{gadX}$-YFP | pSC101 | KanR | 228095 |
| $P_{gadX}$-CFP | $P_{recA}$-YFP | pSC101 | KanR | 228094 |
| $P_{gadX}$-CFP | $P_{const}$-YFP | pSC101 | KanR | 228096 |
| $P_{recA}$-CFP | $P_{gadX}$-YFP | pSC101 | KanR | 228098 |
| $P_{recA}$-CFP | $P_{recA}$-YFP | pSC101 | KanR | 228097 |
| $P_{recA}$-CFP | $P_{const}$-YFP | pSC101 | KanR | 228099 |
| $P_{const}$-CFP | $P_{gadX}$-YFP | pSC101 | KanR | 228102 |
| $P_{const}$-CFP | $P_{recA}$-YFP | pSC101 | KanR | 228101 |
| $P_{const}$-CFP | $P_{const}$-YFP | pSC101 | KanR | 228100 |

**Table S2.** Microscopy imaging settings. Different settings were used to keep fluorescence within the camera detection range. Calibration experiments were conducted to correct for these different settings (Figs. S4 and S5).

| $P_x$-CFP / $P_y$-YFP | CFP imaging settings | YFP imaging settings |
|---|---|---|
| $P_{gadX}$-CFP / $P_{gadX}$-YFP | 25%, 70ms | 30%, 350ms |
| $P_{gadX}$-CFP / $P_{recA}$-YFP | 25%, 70ms | 30%, 350ms |
| $P_{gadX}$-CFP / $P_{const}$-YFP | 25%, 70ms | 30%, 350ms |
| $P_{recA}$-CFP / $P_{gadX}$-YFP | 33%, 200ms | 33%, 50ms |
| $P_{recA}$-CFP / $P_{recA}$-YFP | 25%, 70ms | 30%, 350ms |
| $P_{recA}$-CFP / $P_{const}$-YFP | 33%, 200ms | 30%, 350ms |
| $P_{const}$-CFP / $P_{gadX}$-YFP | 25%, 70ms | 33%, 50ms |
| $P_{const}$-CFP / $P_{recA}$-YFP | 25%, 70ms | 30%, 350ms |
| $P_{const}$-CFP / $P_{const}$-YFP | 25%, 70ms | 30%, 350ms |

**Supplementary Movie Captions**

**Movie S1.** Representative example of cells growing in the mother machine microfluidic device with the $P_{gadX}$-CFP / $P_{gadX}$-YFP reporter plasmid. Time is in HH:MM and scale bar is 10 µm. The mother cell in chamber 17 from the left is the cell portrayed in Fig. 2A. Ciprofloxacin addition at hour 10 is shown by the addition of media with red dye in the media flow channel. The videos continue for 18 hours after ciprofloxacin addition to display cells that survived and died after antibiotic exposure.

**Movie S2.** Representative example of cells growing in the mother machine microfluidic device with the $P_{gadX}$-CFP / $P_{recA}$-YFP reporter plasmid. Time is in HH:MM and scale bar is 10 µm. The mother cell in chamber 9 from the left is the cell portrayed in Fig. 2A. Ciprofloxacin addition at hour 10 is shown by the addition of media with red dye in the media flow channel. The videos continue for 18 hours after ciprofloxacin addition to display cells that survived and died after antibiotic exposure.

**Movie S3.** Representative example of cells growing in the mother machine microfluidic device with the $P_{gadX}$-CFP / $P_{const}$-YFP reporter plasmid. Time is in HH:MM and scale bar is 10 µm. The mother cell in chamber 13 from the left is the cell portrayed in Fig. 2A. Ciprofloxacin addition at hour 10 is shown by the addition of media with red dye in the media flow channel. The videos continue for 18 hours after ciprofloxacin addition to display cells that survived and died after antibiotic exposure.

**Movie S4.** Representative example of cells growing in the mother machine microfluidic device with the $P_{recA}$-CFP / $P_{gadX}$-YFP reporter plasmid. Time is in HH:MM and scale bar is 10 µm. The mother cell in chamber 9 from the left is the cell portrayed in Fig. 2A. Ciprofloxacin addition at hour 10 is shown by the addition of media with red dye in the media flow channel. The videos continue for 18 hours after ciprofloxacin addition to display cells that survived and died after antibiotic exposure.

**Movie S5.** Representative example of cells growing in the mother machine microfluidic device with the $P_{recA}$-CFP / $P_{recA}$-YFP reporter plasmid. Time is in HH:MM and scale bar is 10 µm. The mother cell in chamber 3 from the left is the cell portrayed in Fig. 2A. Ciprofloxacin addition at hour 10 is shown by the addition of media with red dye in the media flow channel. The videos continue for 18 hours after ciprofloxacin addition to display cells that survived and died after antibiotic exposure.

**Movie S6.** Representative example of cells growing in the mother machine microfluidic device with the $P_{recA}$-CFP / $P_{const}$-YFP reporter plasmid. Time is in HH:MM and scale bar is 10 µm. The mother cell in chamber 15 from the left is the cell portrayed in Fig. 2A. Ciprofloxacin addition at hour 10 is shown by the addition of media with red dye in the media flow channel. The videos continue for 18 hours after ciprofloxacin addition to display cells that survived and died after antibiotic exposure.

**Movie S7.** Representative example of cells growing in the mother machine microfluidic device with the $P_{const}$-CFP / $P_{gadX}$-YFP reporter plasmid. Time is in HH:MM and scale bar is 10 µm. The mother cell in chamber 11 from the left is the cell portrayed in Fig. 2A. Ciprofloxacin addition at hour 10 is shown by the addition of media with red dye in the media flow channel. The videos continue for 18 hours after ciprofloxacin addition to display cells that survived and died after antibiotic exposure.

**Movie S8.** Representative example of cells growing in the mother machine microfluidic device with the $P_{const}$-CFP / $P_{recA}$-YFP reporter plasmid. Time is in HH:MM and scale bar is 10 µm. The mother cell in chamber 11 from the left is the cell portrayed in Fig. 2A. Ciprofloxacin addition at hour 10 is shown by the addition of media with red dye in the media flow channel. The videos continue for 18 hours after ciprofloxacin addition to display cells that survived and died after antibiotic exposure.

**Movie S9.** Representative example of cells growing in the mother machine microfluidic device with the $P_{const}$-CFP / $P_{const}$-YFP reporter plasmid. Time is in HH:MM and scale bar is 10 µm. The mother cell in chamber 10 from the left is the cell portrayed in Fig. 2A. Ciprofloxacin addition at hour 10 is shown by the addition of media with red dye in the media flow channel. The videos continue for 18 hours after ciprofloxacin addition to display cells that survived and died after antibiotic exposure.
